# Bepridil is potent against SARS-CoV-2 In Vitro

**DOI:** 10.1101/2020.05.23.112235

**Authors:** Erol C. Vatansever, Kai Yang, Kaci C. Kratch, Aleksandra Drelich, Chia-Chuan Cho, Drake M. Mellott, Shiqing Xu, Chien-Te K. Tseng, Wenshe Ray Liu

## Abstract

Guided by a computational docking analysis, about 30 FDA/EMA-approved small molecule medicines were characterized on their inhibition of the SARS-CoV-2 main protease (M^Pro^). Of these tested small molecule medicines, six displayed an IC_50_ value in inhibiting M^Pro^ below 100 μM. Three medicines pimozide, ebastine, and bepridil are basic small molecules. Their uses in COVID-19 patients potentiate dual functions by both raising endosomal pH to slow SARS-CoV-2 entry into the human cell host and inhibiting M^Pro^ in infected cells. A live virus-based microneutralization assay showed that bepridil inhibited cytopathogenic effect induced by SARS-CoV-2 in Vero E6 cells completely at and dose-dependently below 5 μM and in A549 cells completely at and dose-dependently below 6.25 μM. Therefore, the current study urges serious considerations of using bepridil in COVID-19 clinical tests.

## INTRODUCTION

The current worldwide impact of the COVID-19 pandemic has been so profound that it is often compared to that of 1918 influenza pandemic.(1, 2) As of June 10th, 2020, the total global COVID-19 cases had surpassed 7.1 million, among which more than 400,000 had succumbed to death.(3) A modelling study has predicted that this pandemic will continue to affect everyday life and the circumstances may require societies to follow social distancing until 2022.(4) Finding timely treatment options is of tremendous importance to alleviate catastrophic damages of COVID-19. However, the short time window that is required to contain the disease is extremely challenging to a conventional drug discovery process that requires typically many years to finalize a drug and therefore might not achieve its goal before the pandemic ceases. In this January, we did a comparative biochemical analysis between severe acute respiratory syndrome-coronavirus 2 (SARS-CoV-2), the virus that has caused COVID-19, and SARS-CoV-1 that led to an epidemic in China in 2003 and proposed that remdesivir was a viable choice for the treatment of COVID-19.(5) We were excited to see that remdesivir was finally approved for emergency use in the United States and for use in Japan for people with severe symptoms. With only one medicine in stock right now, the virus may easily evade it, leading us once again with no medicine to use. Given the rapid spread and the high fatality of COVID-19, finding alternative medicines is imperative. Drug repurposing stands out as an attractive option in the current situation. If an approved drug can be identified to treat COVID-19, it can be quickly proceeded to clinical trials and manufactured at a large scale using its existing GMP lines. Previously, encouraging results were obtained from repurposing small molecule medicines including teicoplanin, ivermectin, itraconazole, and nitazoxanide.(6-9) These antimicrobial agents were found effective against virus infections.(10) However, a common drawback of all these repurposed drugs is their low efficacy level. One way to circumvent this problem is to combine multiple existing medicines to accrue a synergistic effect. To be able to discover such combinations, breaking down the druggable targets of the SARS-CoV-2 to identify drugs that do not cross-act on each other’s targets is a promising strategy. For example, a recent study showed that triple combination of interferon β-1b, lopinavir-ritonavir, and ribavirin was safe and superior to lopinavir-ritonavir alone for treating COVID-19 patients.(11)

In our January paper, we recommended four SARS-CoV-2 essential proteins including Spike, RNA-dependent RNA polymerase, the main protease (M^Pro^), and papain-like protease as drug targets for the development of anti-COVID-19 therapeutics. Among these four proteins, M^Pro^ that was previously called 3C-like protease (3CL^pro^) provides the most facile opportunity for drug repurposing owing to the ease of its biochemical assays. M^Pro^ is a cysteine protease that processes itself and then cleaves a number of nonstructural viral proteins from two polypeptide translates that are made from the virus RNA in the human cell host.(12) Its relatively large active site pocket and a highly nucleophilic, catalytic cysteine residue make it likely to be inhibited by a host of existing and investigational drugs. Previous work has disclosed some existing drugs that inhibit M^Pro^.(13) However, complete characterization of existing drugs on the inhibition of M^Pro^ is not yet available. Since the release of the first M^Pro^ crystal structure, many computational studies have been carried out to screen existing drugs in their inhibition of M^Pro^ and many potent leads have been proposed.(14-17) However, most of these lead drugs are yet to be confirmed experimentally. To investigate whether some existing drugs can potently inhibit M^Pro^, we have docked a group of selected FDA/EMA-approved small molecule medicines to the active site of M^Pro^ and selected about 30 hit drugs to characterize their inhibition on M^Pro^ experimentally. Our results revealed that a number of FDA/EMA-approved small molecule medicines have high potency in inhibiting M^Pro^ and bepridil completely inhibits cytopathogenic effect (CPE) induced by the SARS-CoV-2 virus in Vero E6 cells at 5 μM and A549 cells at 6.25 μM. Therefore, the current study encourages the clinical use of bepridil in fighting COVID-19.

## RESULTS & DISCUSSION

Deng *et al*. released the first crystal structure of M^Pro^ on Feb 5th, 2020.(13) We chose this structure (the pdb entry 6lu7) as the basis for our initial docking study. M^Pro^ has a very large active site that consists of several smaller pockets for the recognition of amino acid residues in its protein substrates. Three pockets that bind the P1, P2, and P4 residues in a protein substrate potentially interact with aromatic and large hydrophobic moieties.(18) Although the P1’ residue in a protein substrate is a small residue such as glycine or serine, previous studies based on the same functional enzyme from SARS-CoV-1 showed that an aromatic moiety can occupy the site that originally bind the P1’ and P2’ residues in a substrate.(19) Based on this analysis of the M^Pro^ structure, we selected 55 FDA/EMA-approved small molecule medicines that have several aromatic or large hydrophobic moieties inter-connected and did a docking analysis of their binding to M^Pro^. Some of the small molecule medicines used in our docking study were previously reported in other computational studies.(14-17) Autodock was the program we adopted for the docking analysis.(20) The covalent ligand and non-bonded small molecules in the structure of 6lu7 was removed to prepare the protein structure for docking. Four residues His41, Met49, Asnl42, and Glnl89 that have shown conformational variations in the SARS-CoV-1 enzyme were set flexible during the docking process. We carried out a genetic algorithm method with 100 runs to dock each small molecule medicine to the enzyme. We collected the lowest binding energy from the total 100 runs for each small molecule medicine and summarized them in Table 1. Among all 55 small molecule drugs that we used in the docking study, 29 showed a binding energy lower than -8.3 kcal/mol. We chose these molecules to do further experimental characterizations.

**Table 1:**
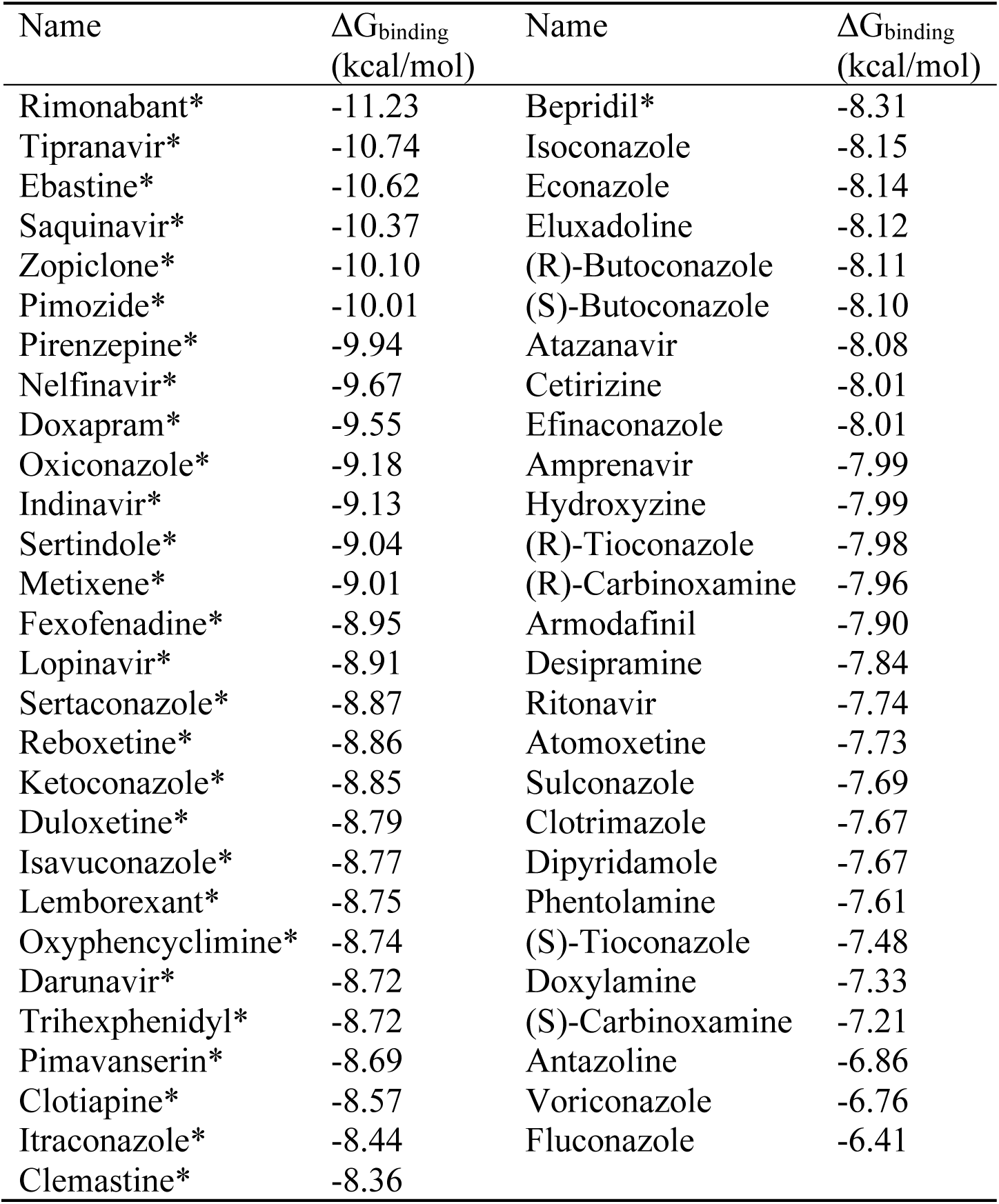
Docking results of small molecule medicines (Compounds whose IC_50_ values were tested are asterisked.)

To express M^Pro^ for experimental characterizations of 29 selected small molecule medicines, we fused the M^Pro^ gene to a superfolder green fluorescent protein (sfGFP) gene and a 6xHis tag at its 5’ and 3’ ends respectively in a pBAD-sfGFP plasmid that we used previously in the lab. SfGFP is known to stabilize proteins when it is genetically fused to them.(21) We designed a TEV protease cleavage site between sfGFP and M^Pro^ for the TEV-catalyzed proteolytic release of M^Pro^ from sfGFP after we expressed and purified the fusion protein. We placed the 6xHis tag right after the M^Pro^ *C*-terminus. The addition of this tag was for straightforward purification with Ni-NTA resins. We expected that the TEV protease cleavage of sfGFP would activate M^Pro^ to cleave the *C*-terminal 6xHis tag so that a finally intact M^Pro^ protein would be obtained. We carried out the expression in *E. coli* TOP10 cells. To our surprise, after expression there was a minimal amount of the fusion protein that we were able to purify. The analysis of the cell lysate showed clearly the cleavage of a substantial amount of M^Pro^ from sfGFP. Since we were not able to enrich the cleaved M^Pro^ using Ni-NTA resins, the *C*-terminal 6xHis tag was apparently cleaved as well. TEV protease is a cysteine protease that cleaves after the Gln residue in the sequence Glu-Asn-Leu-Tyr-Phe-Gln-(Gly/Ser).(22) M^Pro^ is known to cleave the sequence Thr-Val-Leu-Gln-(Gly/Ser).(23) The two cleavage sites share a same P1 residue. It was evident in our expression work that M^Pro^ is able to efficiently cleave the TEV protease cutting site to maturate inside *E. coli* cells. It is likely that M^Pro^ has a substrate promiscuity higher than what we have learnt from the SARS-CoV-1 enzyme. To purify the cleaved and maturated M^Pro^, we used ammonium sulfate to precipitate it from the cell lysate and then used the ion exchange and size exclusion chromatography to isolate it to more than 95% purity. We designed and synthesized a fluorogenic coumarin-based hexapeptide substrate (Sub1) and a FRET-based decapeptide substrate (Sub2) and acquired a commercial FRET-based tetradecapeptide substrate (Sub3) (Figure 1A). The test of enzyme activities on the three substrates indicated that the enzyme had low activity toward Sub1 (Figure 1B) and its activity on Sub3 was higher than that on Sub2 (Figure 1C) under our assay conditions. We subsequently used Sub3 in all following inhibition analysis. To identify an optimal enzyme concentration for use in our inhibition analysis, we tested activities of different concentrations of M^Pro^ on 10 µM Sub3, the detected catalytic rate of the Sub3 cleavage was not proportional to the enzyme concentration (Figure 1D). When the enzyme concentration decreased from 50 nM to 10 nM, the Sub3 cleavage rate dropped roughly proportionally to the square of the concentration decrease, characteristics of second-order kinetics. This observation supports previous claims that the enzyme needs to dimerize in order to be active.(24) In all the following assays, 50 nM M^Pro^ and 10 µM Sub3 were used throughout.

**Figure 1:**
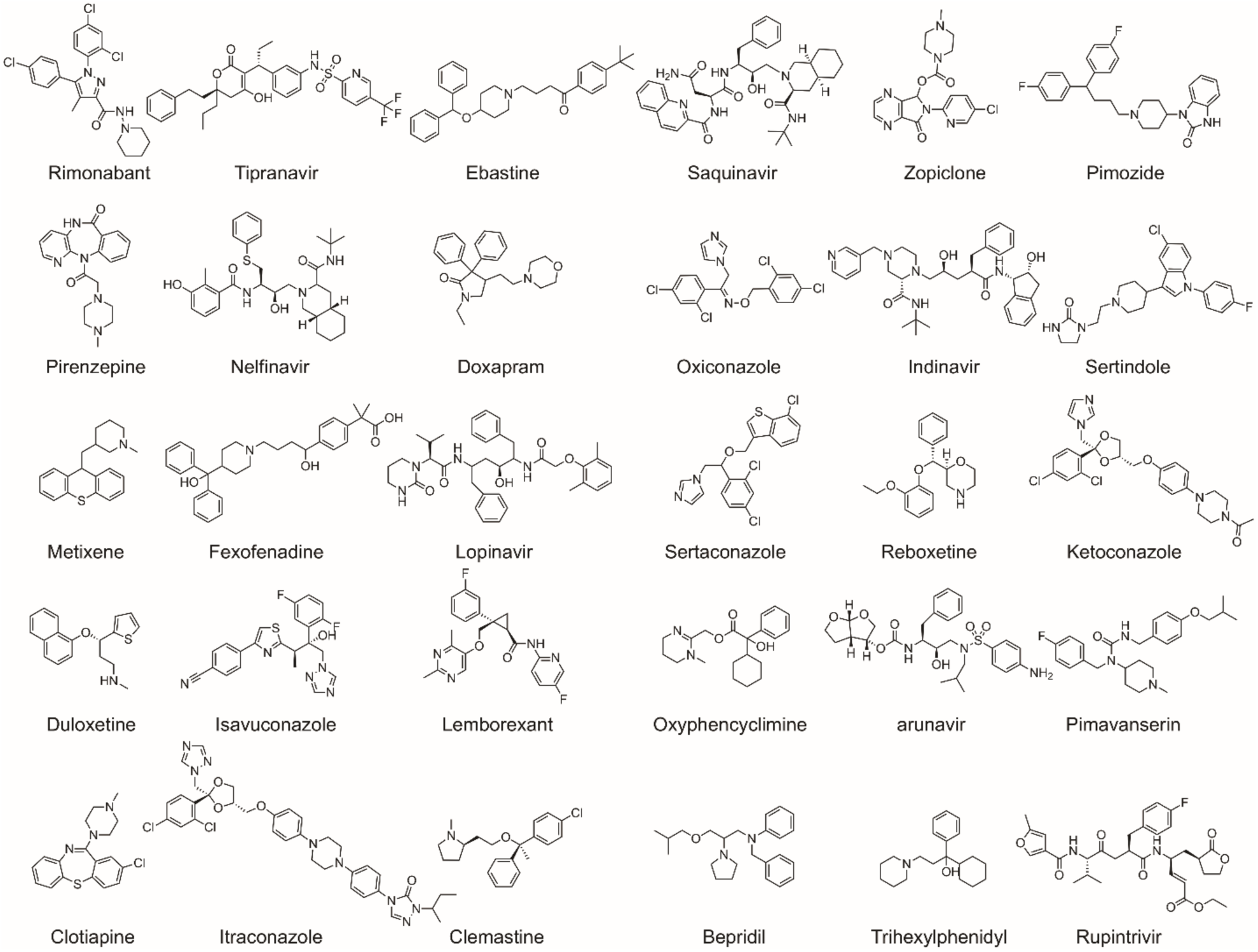
Structures of 29 FDA/EMA-approved medicines and rupintrivir whose IC_50_ values in inhibiting M^Pro^ were determined in the study.

We purchased all 29 small molecule medicines from commercial providers without further purification and characterization. Rupintrivir is a previously developed 3C protease inhibitor.(25) It has a key lactone side chain in the P1 residue that has a demonstrated role in tight binding to 3C and 3CL proteases. Since it has been an investigational antiviral medicine, we purchased it as well with a hope that it could be a potent inhibitor of M^Pro^. We dissolved most purchased small molecule medicines in DMSO to make 5 mM stock solutions and proceeded to use these stock solutions to test inhibition on M^Pro^. Except itraconazole that has low solubility in DMSO, all tested small molecule medicines were diluted to a 1 mM final concentration in the inhibition assay conditions. We maintained 20% DMSO in the final assay condition to prevent small molecule medicines from precipitation. The activity of M^Pro^ in 20% DMSO was a little lower than that in a regular buffer but satisfied our assay requirement. An M^Pro^ activity assay in the absence of a small molecule medicine was set up as a comparison. Triplicate repeats were carried out for all tested small molecules and the control. The results presented in Figure 2 displayed two easily discernable characteristics. First, about half of the tested compounds showed strong inhibition of M^Pro^ at the 1 mM concentration level (itraconazole was at 0.14 mM due to its low solubility in DMSO), supporting the practical use of a docking method in guiding the drug repurposing research of COVID-19. Second, several small molecule medicines including fexofenadine, indinavir, pirenzepine, reboxetine, and doxapram clearly activated M^Pro^ (> 15%). This was to the contrary of what the docking program predicted. This observation strongly suggests that frontline clinicians need to exhibit caution in repurposing medicines for COVID-19 patients before they are thoroughly investigated on influencing the SARS-CoV-2 biology. A not-well-understood drug might deteriorate the already devastating symptoms in COVID-19 patients. Although it is not the focus of the current study, the observation that M^Pro^ can be activated by existing drugs needs to be further investigated.

**Figure 2:**
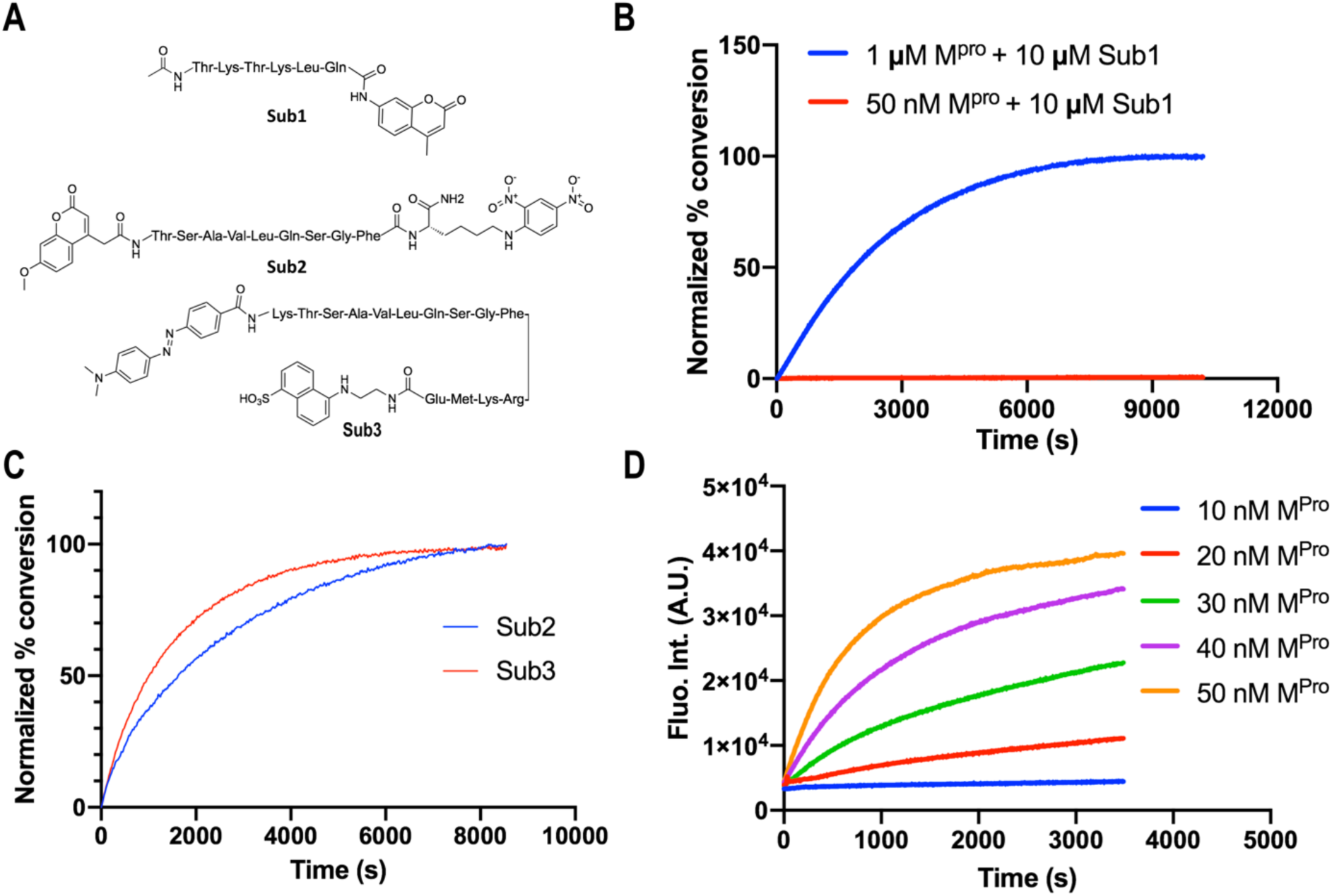
Activity of M^Pro^. (A) The structures of three substrates. (B) Activity of 50 nM M^Pro^ on 10 μM Sub1. (C) Activity of 50 nM M^Pro^ on 10 μM Sub2 and Sub3. The florescence signals are normalized for easy comparison. (D) Activity of different concentrations of M^Pro^ on 10 μM Sub3.

We selected 17 small molecule medicines and rupintrivir that displayed strong inhibition of M^Pro^ to conduct further characterizations of their IC_50_ values in inhibiting M^Pro^ by varying the small molecule concentration from 1 µM to 10 mM. Results collectively presented in Figure 3 identifies that of the 18 tested compounds, 7 had an IC_50_ value below 100 µM. These include pimozide, ebastine, rupintrivir, bepridil, sertaconazole, rimonabant, and oxiconazole. Pimozide, ebastine, and bepridil were the three most potent FDA/EMA-approved medicines with IC_50_ values as 42 ± 2, 57 ± 12 and 72 ± 12 µM, respectively. Although rupintrivir is a covalent inhibitor that was developed specifically for 3C and 3CL proteases, its IC_50_ value (68 ± 7 µM) is higher than that of pimozide and ebastine. The relatively low activity of rupintrivir in inhibiting M^Pro^ might be due to the change of the amide bond between the P2 and P3 residues to an methyleneketone. This conversion served to increase the serum stability of rupintrivir, but has likely eliminated a key hydrogen bonding interaction with M^Pro^.(13) The repurposing of HIV medicines for the treatment of COVID-19, particularly those targeting HIV1 protease, has been area of much attention. In fact, the cocktail of lopinavir and ritonavir was previously tested in China for the treatment of COVID-19.(26) The IC_50_ value of lopinavir in inhibiting M^Pro^ is about 500 µM, which possibly explains why this anti-HIV viral cocktail demonstrated no significant benefit for treating patients. Nelfinavir was previously shown having high potency in inhibiting the entry of SARS-CoV-2 into mammalian cell hosts. A cell-based study in inhibiting the SARS-CoV-2 entry indicated a 1 µM EC_50_ value.(27, 28) However, our IC_50_ determination against M^Pro^ resulted in a value of 234 ± 5 µM, highlighting that nelfinavir likely inhibits another key SARS-CoV-2 enzyme or protein or interferes with key cellular processes that are required for the SARS-CoV-2 entry into host cells. These possibilities need to be studied further. Structurally the two most potent medicines pimozide and ebastine share a same diphenylmethyl moiety. A spatially similar structure moiety N-phenyl-N-benzylamine exists in bepridil. Our docking results suggested a same binding mode for this similar structure moiety in all three drugs (Figure 5). The two aromatic rings occupy the enzyme pockets that associate with the P2 and P4 residues in a substrate. This observation is in line with a crystallographic study that showed two aromatic rings with a single methylene linker bound to the active site of the SARS-CoV-1 enzyme.(29) We believe that the inclusion of the diphenylmethyl moiety in structure-activity relationship studies of M^Pro^-targeting ligands will likely contribute to the identification of both potent and high cell-permeable M^Pro^ inhibitors. Figure 4 also revealed large variations in Hill coefficients of IC_50_ curves for different small molecule medicines (IC_50_ values and Hill coefficients are summarized in Table 2). Duloxetine and zopiclone gave the two highest Hill coefficients with a gradual M^Pro^ activity decrease over an increasing inhibitor concentration. On the contrary, saquinavir and lopinavir yielded lowest Hill coefficients with highly steep IC_50_ curves. There are three possible explanations for the large discrepancy in Hill coefficients. It could be attributed to different solubility of tested compounds. It is possible that a high DMSO percentage and a relatively high inhibitor concentration created phase transition for some inhibitors.(30) A high Hill coefficient may also be due to different ligand to enzyme ratios when tested compounds bind to M^Pro^. An additionally possible reason is the co-existence of the M^Pro^ dimer and monomer in the assay conditions. A previous report showed a K_d_ value of the M^Pro^ dimerization as 2.5 µM.(18) In theory, the catalytically active dimer species was present at a very low concentration in our assay conditions, leaving the catalytically inactive monomer species as the major form of M^Pro^. In this situation, the inhibitors that preferentially bind to the M^Pro^ dimer and the inhibitors that have a higher affinity to the M^Pro^ monomer might yield different Hill coefficients.

**Table 2:** IC_50_ and Hill coefficient values of 18 identified inhibitors

| Name | IC<sub>50</sub> (μM) | Hill Slope |
| --- | --- | --- |
| Pimozide | 42 ± 2 | 3.1 ± 0.4 |
| Ebastine | 57 ± 12 | 1.5 ± 0.2 |
| Rupintrivir | 68 ± 7 | 1.4 ± 0.2 |
| Bepridil | 72 ± 3 | 2.9 ± 1.0 |
| Sertaconazole | 76 ± 2 | 3.5 ± 0.2 |
| Rimonabant | 85 ± 3 | 5.0 ± 0.4 |
| Oxiconazole | 99 ± 6 | 3.8 ± 0.4 |
| Itraconazole | 111 ± 35 | 1.6 ± 0.2 |
| Tipranavir | 180 ± 20 | 1.4 ± 0.2 |
| Nelfinavir | 234 ± 15 | 5.4 ± 1.0 |
| Zopiclone | 349 ± 77 | 1.2 ± 0.2 |
| Trihexyphenidyl | 370 ± 53 | 8.9 ± 6.4 |
| Saquinavir | 411 ± 6 | 26.8 ± 2.6 |
| Isavuconazole | 438 ± 11 | 5.2 ± 0.7 |
| Lopinavir | 486 ± 2 | 29.9 ± 2.4 |
| Clemastine | 497 ± 148 | 11.2 ± 7.3 |
| Metixene | 635 ± 43 | 8.7 ± 5 |
| Duloxetine | 3047 ± 634 | 0.93 ± 0.07 |

**Figure 3:**
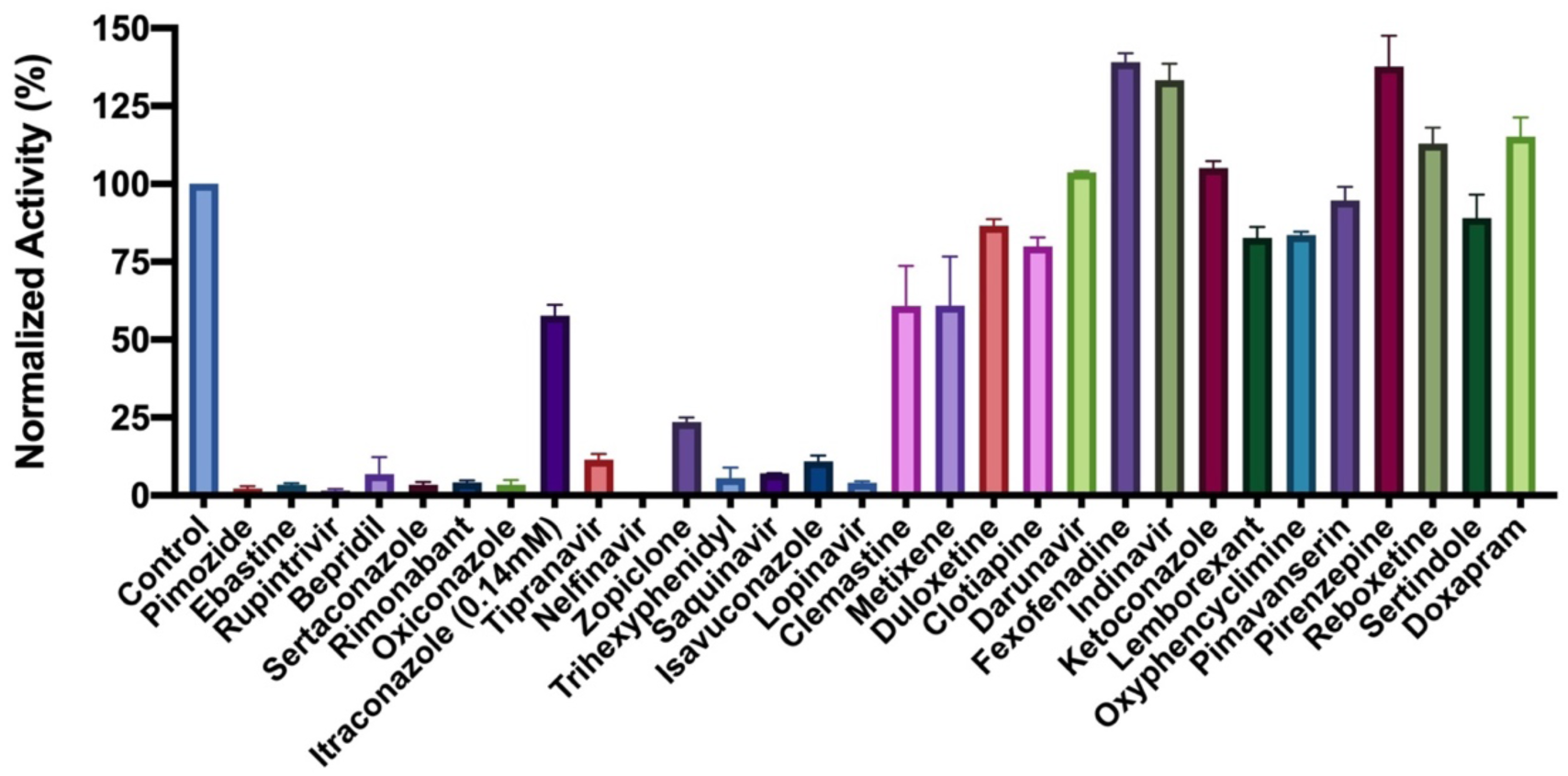
Initial screening of M^pro^ inhibition by 29 FDA/EMA-approved medicines and rupintrivir. 1 mM (0.14 mM for Itraconazole due to its low solubility in DMSO) was used for each inhibitor to perform the inhibition assay. Fluorescence intensity was normalized with respect to the control that had no small molecule provided. Triplicate experiments were performed for each compound, and the value was presented as mean ± standard error (SE).

**Figure 4:**
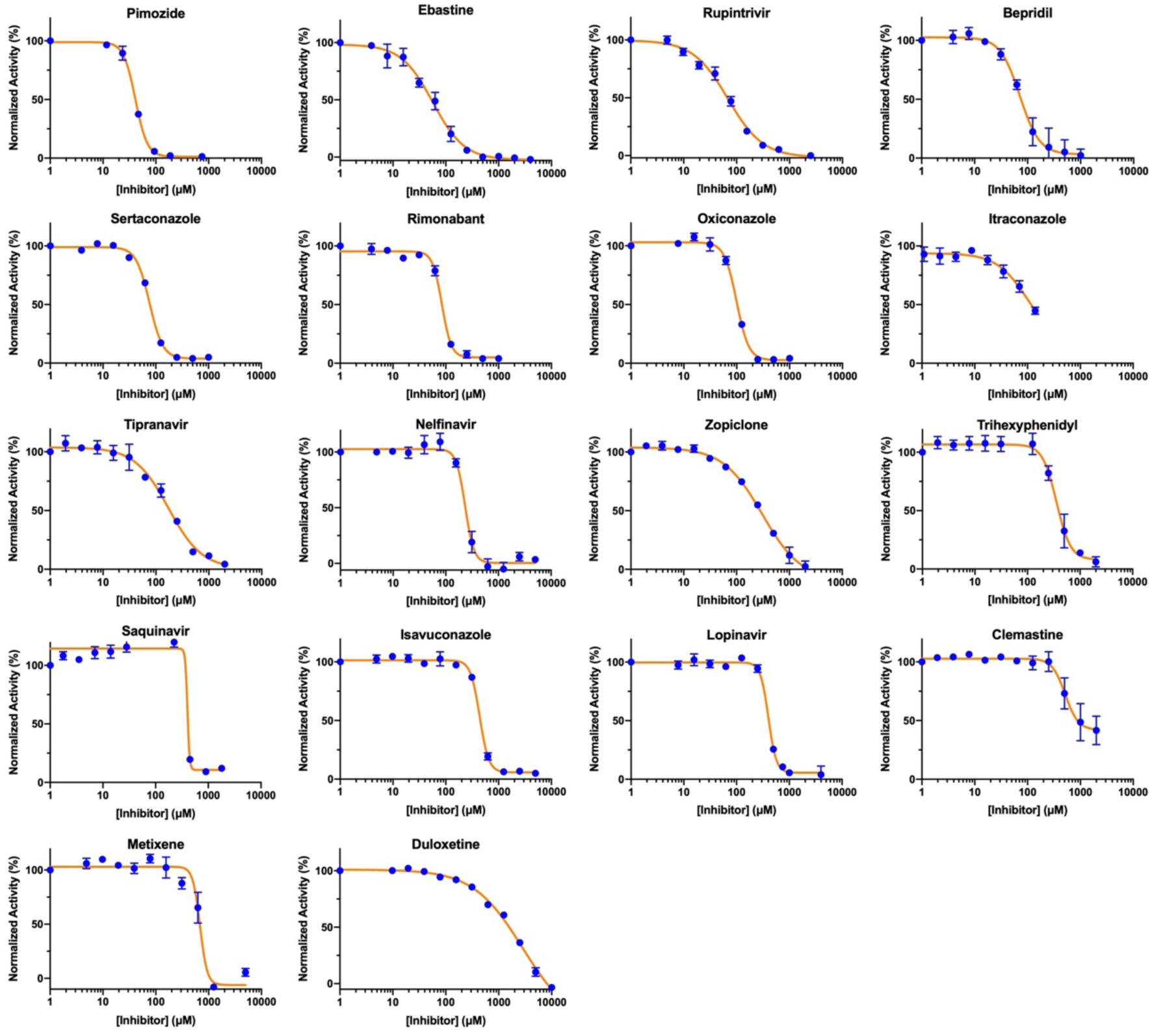
IC_50_ assays for 18 small molecule medicines on their inhibition of M^pro^. Triplicate experiments were performed for each compound, and the IC_50_ value was presented as mean ± standard error (SE). GraphPad Prism 8.0 was used to perform data analysis.

**Figure 5:**
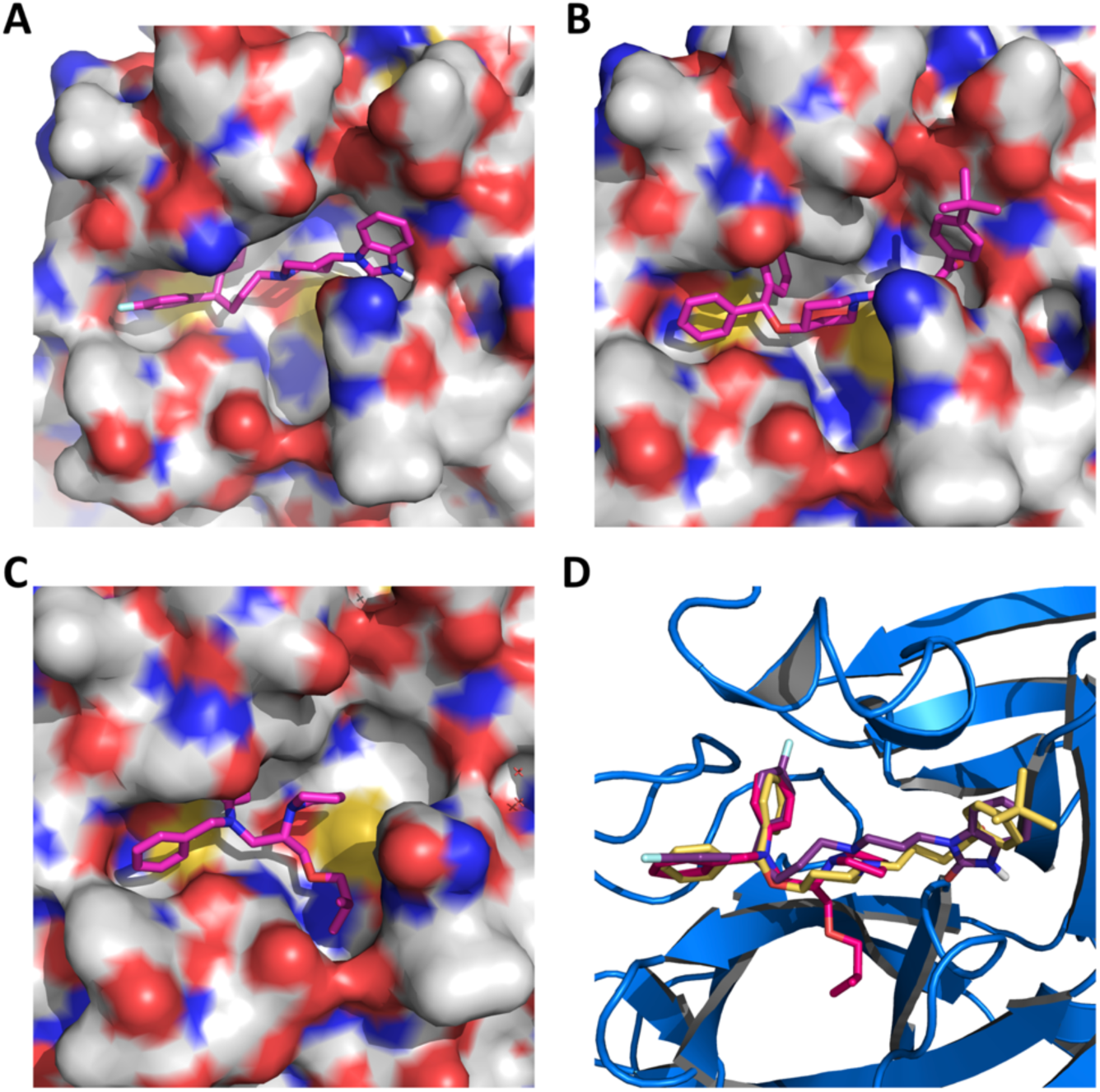
Pimozide (A), ebastine (B), bepridil (C), and their overlay (D) in the active site of M^Pro^. The protein surface topography in A, B, and C is presented to show the concaved active site.

From cell biology point of view, our three lead compounds share similarities with some proposed COVID-19 treatment options. There are reports on the investigation of using hydroxychloroquine to treat COVID-19 patients.(31, 32) A likely mechanism of action for hydroxychloroquine is its ability to raise endosomal pH that impacts significantly activities of endosomal proteases that may be required to process the virus membrane proteins.(33, 34) Our top three hits pimozide, ebastine, and bepridil are all basic small molecules that can potentiate a similar effect.(35) Among the three drugs, bepridil can be very interesting because it previously provided 100% protection from Ebola virus infections in mice at a dose of 12 mg/kg.(36) Bepridil is a calcium channel blocker with a significant anti-anginal activity. For patients with chronic stable angina, recommended daily dose of bepridil is 200-400 mg.(29) Mice administered with a bepridil dose as high as 300 mg/kg/day did not show alteration in mating behavior and reproductive performance, indicating that bepridil has very low toxicity.(37) Moreover, a previous study showed that bepridil can increase the pH of acidic endosomes.(38) Administration of a high dose of bepridil may have dual functions to slow down the virus replication in host cells by both inhibiting M^Pro^ and raising the pH of endosomes. To demonstrate this prospect, we conducted a live virus-based microneutralization assay to evaluate efficacy of pimozide, ebastine and bepridil in their inhibition of SARS-CoV-2 infection in Vero E6 cells. Vero E6 is a cell line isolated kidney epithelial cells from African Green Monkey. We tested three medicines in a concentration range from 0.16 to 200 μM. Cytopathogenic effect (CPE) was clearly observable for pimozide and ebastine at all tested concentrations. However, bepridil prevented completely the SARS-CoV-2-induced CPE in Vero E6 cells when the concentration reached 5 μM and inhibited CPE in a dose dependent manner below 5 μM (Table 3A). It did not display cellular toxicity until the concentration reached 50 μM. A parallel test in A549 cells that were derived from human alveolar epithelial cells showed that bepridil prevented SARS-CoV-2-induced CPE completely at 6.25 μM and inhibited CPE in a dose dependent manner below 6.25 μM but did not display a cytotoxic effect when the concentration reached 200 μM (Table 3B). The complete prevention of SARS-CoV-2-induced CPE in Vero E6 and A549 cells by bepridil at a concentration much lower than its IC_50_ value for inhibiting M^Pro^ is likely due to the aforementioned dual functions or other cellular effects of bepridil. In patients, bepridil can reach a state C_max_ as 3.72 μM.(39) This concentration is effective in inhibiting SARS-CoV-2 based on our virus microneutralization analysis. Collectively, our results indicate that bepridil is effective in preventing SARS-CoV-2 from entry and replication in mammalian cell hosts. Therefore, we urge clinical tests of bepridil in the treatment of COVID-19.

**Table 3:** SARS-CoV-2 induced CPE in (A) Vero E6 and (B) ACE2-expressing A549 cells in the presence of bepridil

| Bepridil (μM) | Repeat 1 | Repeat 2 |
| --- | --- | --- |
| 200 | C<sup>a</sup> | C |
| 100 | C | C |
| 50 | C | C |
| 20 | ND<sup>b</sup> | ND |
| 10 | ND | ND |
| 5 | ND | ND |
| 2.5 | CPE | CPE |
| 1.25 | CPE | CPE |
| 0.62 | CPE | CPE |
| 0.31 | CPE | CPE |
| 0.16 | CPE | CPE |

**B.**
| Bepridil (μM) | Repeat 1 | Repeat 2 |
| --- | --- | --- |
| 200 | ND | ND |
| 100 | ND | ND |
| 50 | ND | ND |
| 25 | ND | ND |
| 12.5 | ND | ND |
| 6.25 | ND | ND |
| 3.12 | CPE | CPE |
| 1.56 | CPE | CPE |

<sup>a</sup>Cytotoxicity; <sup>b</sup>Both cytotoxicity and CPE are not detected.

## CONCLUSION

Guided by a computational docking analysis, we experimentally characterized about 30 FDA/EMA-approved drugs on their inhibition of the essential M^Pro^ enzyme of the COVID-19 pathogen SARS-CoV-2. From the study, we identified six FDA/EMA-approved drugs that can potently inhibit M^Pro^ with an IC_50_ value lower than 100 µM. One medicine bepridil exhibited strong inhibition of SARS-CoV-2 from entry and replication inside Vero E6 cells at a low micromolar concentration. Given that bepridil has been previously explored to treat Ebola infected patients, we urge a serious consideration of its clinical tests in treating COVID-19. Our current study indicates that there is a large amount of FDA/EMA-approved drug space open for exploration that could hold promise for repurposing existing drugs to target COVID-19. Performing screening studies on different SARS-CoV-2 protein targets are necessary to uncover existing medicines that may be combined for cocktail treatments of COVID-19. More explorations in this direction are imperative.

## MATERIALS AND METHODS

### Chemicals

We purchased econazole nitrate, duloxetine hydrochloride, doxapram hydrochloride monohydrate, clemastine fumarate salt, sertaconazole nitrate, isavuconazole, rupintrivir, and zopiclone from Sigma-Aldrich, pimavanserin, trihexyphenidyl hydrochloride, reboxetine mesylate, sertindole, bepridil hydrochloride, darunavir, nelfinavir mesylate, indinavir sulfate, lopinavir, tipranavir, saquinavir, pirenzepine hydrochloride, oxiconazole nitrate, pimozide, and rimonabant from Cayman Chemicals, ebastine and itraconazole from Alfa Aesar, metixene hydrochloride hydrate and lemborexant from MedChemExpress, fexofenadine hydrochloride from TCI Chemicals, ketoconazole from Acros Organics, clotiapine from Enzo Life Sciences, and oxyphencyclimine from Boc Sciences. We acquired Sub3 with the sequence as DABCYL-Lys-Thr-Ser-Ala-Val-Leu-Gln-Ser-Gly-Phe-Arg-Lys-Met-Glu-EDANS from Bachem Inc.

### Docking

Autodock 4 was used for all docking analysis. For each small molecule, the genetic algorithm-based calculation was carried out for 100 runs with each run having a maximal number of evaluations as 2,500,000.

### M^pro^ Expression and Purification

We constructed the plasmid pBAD-sfGFP-M^pro^ from pBAD-sfGFP. The M^Pro^ gene was inserted between DNA sequences that coded sfGFP and 6xHis. The overall sfGFP-M^Pro^-6xHis fusion gene was under control of a pBAD promoter. The antibiotic selection marker was ampicillin. To express sfGFP-M^Pro^-6xHis, E. coli TOP10 cells were transformed with pBAD-sfGFP-M^Pro^. A single colony was picked and grew in 5 mL LB medium with 100 µg/mL ampicillin overnight. In the next day, we inoculated this starting culture into 5 L 2xYT medium with 100 µg/mL ampicillin in 5 separate flasks at 37 °C. When the OD reached to 0.6, we added L-arabinose (working concentration as 0.2%) to each flask to induce protein expression at 37 °C for 4 h. Then, the cells were pelleted at 4000 rpm at 4 °C, washed with cold PBS and stored in -80 °C until purification. To purify the expressed protein, we re-suspended frozen cells in a 125 mL buffer containing Tris pH 7.5, 2.5 mM DTT, and 1.25 mg lysozyme. We sonicated resuspended cells using a Branson 250W sonicator with 1 second on, 4 second off, and a total 5 min 60% power output in two rounds. After sonication, we spun down the cellular debris at 16000 rpm for 30 min at 4 °C. We collected the supernatant and recorded the volume. The whole cell lysate analysis showed almost all of the fusion protein was hydrolyzed to two separate proteins sfGFP and M^Pro^. We were able to obtain an insignificant amount of M^Pro^ when Ni-NTA resins were used for purification. Therefore, we did ammonium sulfate precipitation of the whole cell lysate method. This was done by the addition of a saturated ammonium sulfate solution at 0 °C. We collected the fraction between 30% and 40% of ammonium sulfate. We dissolved the collected fraction in buffer A (20 mM Tris, 10 mM NaCl, and 1 mM DTT at pH 8.0) and dialyzed the obtained solution against the same buffer to remove ammonium sulfate. Then, we subjected this solution to anion exchange column chromatography using Q sepharose resins. We eluted proteins from the Q sepharose column by applying a gradient with increasing the concentration of buffer B (20 mM Tris, 1 M NaCl, and 1 mM DTT at pH 8.0). We concentrated the eluted fractions that contained M^Pro^ and subject the concentered solution to size exclusion chromatography using a HiPrep 16/60 Sephacryl S-100 HR column with a mobile phase containing 10 mM sodium phosphate, 10 mM NaCl, 0.5 mM EDTA and 1 mM DTT at pH 7.8. The final yield of the purified enzyme was 1 mg/L with respect to the original expression medium volume. We determined the concentration of the finally purified M^pro^ using the Pierce(tm) 660nm protein assay and aliquoted 10 µM M^Pro^ in the size exclusion chromatography buffer for storage in -80 °C.

### The synthesis of Sub1

We loaded the first amino acid (0.5 mmol, 2 equiv.) manually on chlorotrityl chloride resin (0.52 mmol/g loading) on a 0.25 mmol scale by the addition of DIPEA (3 equiv.). After addition of the first amino acid, automated Fmoc-based solid phases synthesis was performed using a Liberty Blue automated peptide synthesizer. Deprotection of the Fmoc group was carried out with 20% piperidine/DMF. Coupling was done with a Fmoc-protected amino acid (0.75 mmol, 3.0 equiv.) and the coupling reagent HATU (0.9 mmol, 3.6 equiv.) and DIEA in NMP (1 mmol, 4.0 equiv.). The final amino acid was capped by the addition of 25% acetic anhydride (v/v) in DMF and DIEA (0.2mmol, 2.0 equiv.). Coumarin coupling was performed in anhydrous THF using T3P in EtOAc (50% w/v) (3.0 equiv.), DIEA (3 equiv.) and 7-amino-4methyl-coumarin (0.8 equiv.) and mixed for 16 h. The solvent was removed and the peptide was dissolved in DCM and washed with H_2_O (4x) followed by HCl (2x) and brine (1x). The organic layer was dried with Na_2_SO_4_, filtered and concentrated. Global deprotection was then carried out using triisopropylsilane (5%) and trifluoroacetic acid (30%) v/v in DCM and mixed for 2-3 h to result in the crude substrate. The peptide was then purified by semi-preparative HPLC and the fractions containing pure product were pooled, concentrated, and analyzed by LC-MS for purity.

### The synthesis of Sub2

We performed automated Fmoc-based solid phase synthesis on a Liberty Blue automated peptide synthesizer. Synthesis was conducted on a 0.1 mmol scale with Fmoc Rink amide MBHA resin (0.52 mmol/g loading) and 3 equiv. of protected amino acids. Deprotection of the Fmoc group was carried out with 20% piperidine/DMF. Coupling was done using the desired Fmoc-protected amino acid (0.2 mmol, 2.0 equiv.), coupling reagent Oxyma (0.4 mmol, 4.0 equiv.) and DIC (0.4 mmol, 4.0 equiv.). After the final amino acid had been coupled on, the resin was washed trice with DMF and DCM. Cleavage from the resin was performed using trifluoroacetic acid (95%), triisopropylsilane (2.5%) and water (2.5%) with agitation for 4 h. The peptide was drained into cold methyl tert-butyl ether where it precipitated out. We centrifuged the precipitate, decanted mother liquor, dissolved the pellet in DMF and then purified the peptide by LCMS.

### Screening assay

5 mM stock solutions of medicines were prepared in DMSO. The final screening assay conditions contained 50 nM M^Pro^, 10 µM Sub3, and 1 mM medicine. We diluted enzyme stock and substrate stock solutions using a buffer containing 10 mM sodium phosphate, 10 mM NaCl, and 0.5 mM EDTA at pH 7.8 for reaching desired final concentrations. We ran the assay in triplicates. First, we added 30 µL of a 167 nM M^Pro^ solution to each well in a 96-well plate and then provided 20 µL of 5 mM stock solutions of medicines in DMSO. After a brief shaking, we incubated the plate at 37°C for 30 min. Then we added 50 µL of a 20 µM Sub3 solution to initiate the activity analysis. The EDANS fluorescence with excitation at 336 nm and emission at 455 nm from the cleaved substrate was detected. We determined the fluorescence increasing slopes at the initial 5 min and normalized them with respect to the control that had no inhibitor provided.

### Inhibition analysis

The final inhibition assay conditions contained 50 nM M^Pro^, 10 µM Sub3, and a varying concentration of an inhibitor. Similar to screening assay, we diluted enzyme stock and substrate stock solutions using a buffer containing 10 mM sodium phosphate, 10 mM NaCl, and 0.5 mM EDTA at pH 7.8 for reaching desired final concentrations. We ran the assay in triplicates. For the inhibition analysis, we added 30 µL of a 167 nM M^Pro^ solution to each well in a 96-well plate and then provided 20 µL of inhibitor solutions with varying concentrations in DMSO. After a brief shaking, we incubated the plate at 37°C for 30 min. Then we added 50 µL of a 20 µM Sub3 solution to initiate the activity analysis. We monitored the fluorescence signal and processed the initial slopes in the same way described in screening assay part. We used GraphPad 8.0 to analyze the data and used the [Inhibitor] vs. response - Variable slope (four parameters) fitting to determine the values of both IC_50_ and Hill coefficient.

### SARS-CoV-2 inhibition by a cell-based assay

A slightly modified standard live virus-based microneutralization (MN) assay was used as previously described(40-42) to rapidly evaluate the drug efficacy against SARS-CoV-2 infection in Vero E6 and A549 cell cultures. Briefly, confluent Vero E6 and A549 cells grown in 96-wells microtiter plates were pre-treated with serially 2-folds diluted individual drugs in duplicate over eight concentrations for two hours before infection with ∼100 and ∼500, respectively, infectious SARS-CoV-2 particles in 100 µL EMEM supplemented with 2% FBS. Vero E6 and A549 cells treated with parallelly diluted dimethyl sulfoxide (DMSO) with or without virus were included as positive and negative controls, respectively. After cultivation at 37 °C for 4 days, individual wells were observed under the microcopy for the status of virus-induced formation of CPE. The efficacy of individual drugs was calculated and expressed as the lowest concentration capable of completely preventing virus-induced CPE in 100% of the wells. The toxicity to the treated cells was assessed by observing floating cells and altered morphology of adhered Vero E6 and A549 cells in wells under the microcopy. All compounds were dissolved in 100% DMSO as 10 mM stock solutions and diluted in culture media.

## ACKNOWELDGEMENTS

We thank Profs. Thomas Meek and Kevin Burgess for allowing us to use their instruments. This work was supported in part by Texas A&M Presidential Impact Fellowship Fund, National Institutes of Health (grants R01GM127575 and R01GM121584) and Welch Foundation (A-1715).

## AUTHOR CONTRIBUTIONS

W.R.L. conceived the project. E.C.V., K.Y., K.C.K., C.-C.C., A.D., D.M.M., S.X., C.-T.K.T., and W.R.L. designed experiments. E.C.V., K.Y., K.C.K., C.-C.C., A.D., D.M.M. performed the experiments. E.C.V., K.Y. and W.R.L. wrote the manuscript. All authors approved the final manuscript before submission.

